# Rescuing the attentional performance of rats with cholinergic losses by the M1 positive allosteric modulator TAK-071

**DOI:** 10.1101/606343

**Authors:** Aaron Kucinski, Kyra B. Phillips, Ajeesh Koshy Cherian, Martin Sarter

## Abstract

Loss of basal forebrain cholinergic neurons contributes to the severity of the cognitive decline in age-related dementia and to impairments in gait and balance, and the resulting risks for falls, in patients with Parkinson’s disease (PD). Contrasting with the extensive evidence indicating an essential role of cholinergic activity in mediating cognitive, specifically attentional abilities, treatment with conventional acetylcholinesterase inhibitors (AChEIs) has not fulfilled the promise of efficacy of pro-cholinergic treatments. Here we investigated the potential usefulness of a muscarinic M1 positive allosteric modulator (PAM) in an animal model of cholinergic loss-induced impairments in attentional performance. Given evidence indicating that fast, transient cholinergic signaling mediates the detection of cues in attentional contexts, we hypothesized that an M1 PAM amplifies such transient signaling, thereby enhancing and rescuing attentional performance. Rats performed an operant sustained attention task (SAT), including in the presence of a distractor (dSAT) and during a post-distractor (post-dSAT) period assessing their capacity for recovering performance. Basal forebrain infusions of the cholino-specific immunotoxin 192 IgG-saporin impaired SAT performance, and greater cholinergic losses predicted lower post-dSAT performance recovery. Administration of TAK-071 (0.1, 0.3 mg/kg, p.o., administered over 6-day blocks) improved the performance of all rats during the post-dSAT period (main effect of dose). Drug-induced improvement of post-dSAT performance was relatively greater in lesioned rats, irrespective of sex, and also manifested in female control rats. TAK-071 primarily improved perceptual sensitivity (*d’*) in lesioned rats and facilitated the adoption of a more liberal response bias (*B”*_*D*_) in all female rats. Collectively, these findings suggest that TAK-071 may benefit the attentional performance of patients with partial cholinergic losses and specifically in situations that tax top-down, or goal-driven, attentional control.

## INTRODUCTION

The basal forebrain cholinergic system broadly innervates telencephalic regions, including the cortex. Contemporary research has revealed the presence of highly topographic, afferent and efferent projections of basal forebrain neurons, dissociable clusters of basal forebrain cholinergic neurons, target region-specific integration of cholinergic synapses, and the presence of fast cholinergic signaling (scale of seconds) in performing rodents (e.g., Zaborszky, 2002; Parikh et al., 2007; Zaborszky et al., 2008; Howe et al., 2013; Sarter and Kim, 2015; Gritton et al., 2016; Gielow and Zaborszky, 2017; Howe et al., 2017; Yuan et al., 2018; Lean et al., 2019). Together, this evidence suggests that traditional views of this neuronal system as a “diffusely” organized ascending “arousal” system, that relatively slowly (on the scale of minutes) modulates target circuitry, require major revision (see also Hasselmo and Sarter, 2011; Dayan, 2012; Higley and Picciotto, 2014; Ballinger et al., 2016).

Losses of basal forebrain cholinergic neurons have long been associated with the severity of cognitive decline in the age-related dementias (e.g., Mesulam, 2004; Schmitz et al., 2018). Early and still relatively limited cholinergic losses are increasingly considered a biomarker that predicts vulnerability and severity of later cognitive decline (e.g., Schmitz et al., 2016; Cantero et al., 2017; Albin et al., 2018; Schulz et al., 2018). Such early cholinergic losses are specifically associated with the manifestation of impairments in attention and associated deficits, including cognitive-motor interactions (Kim et al., 2017b, a; Kucinski et al., 2019). Assuming that residual cholinergic neurons preserve a degree of cholinergic signaling, as was suggested by evidence from rats with partial losses (e.g., Fadel et al., 1996), and that post-synaptic mechanisms likewise remain functional (Rossner et al., 1995b), pro-cholinergic treatments would be expected to benefit the cognitive and cognitive-motor status of patients with mild cognitive impairments (MCI), early Alzheimer’s disease, and with Parkinson’s disease (PD) who also exhibit cholinergic losses and thus a high propensity for falls (Bohnen and Albin, 2009; Bohnen et al., 2009).

Clearly, however, this prediction has not been convincingly supported by the clinical efficacy of AChEIs (Courtney et al., 2004; Ritchie et al., 2004; Hansen et al., 2008; Maher-Edwards et al., 2011). As the striking and lasting increases in extracellular ACh levels that result from inhibition of AChE appear unlikely to enhance phasic cholinergic pre- and postsynaptic signaling, this class of drugs may not be suitable to assess the potential of pro-cholinergic treatments (see also the discussion in Sarter, 2015). In contrast, the mechanisms exploited by PAMs at post-synaptic muscarinic receptors appear more likely to enhance and perhaps rescue diminished cholinergic signaling (see also Shirey et al., 2009; Digby et al., 2012; Moran et al., 2018; Uslaner et al., 2018). Here we investigated the M1 PAM TAK-071, in part because of the relative selectivity of this drug for acting in the brain and preclinical evidence indicating pro-cholinergic and behavioral efficacy (Sako et al., 2018; Kurimoto et al., 2019).

The contrast between the limited clinical efficacy of AChEIs and strong predictions of such efficacy based on evidence from conventional rodent models and behavioral tasks has sustained long-standing concerns about the usefulness of preclinical approaches to characterize pro-cognitive, pro-cholinergic treatments (Sarter et al., 1992b, a; Sarter, 2004; Sarter and Tricklebank, 2012). Here we investigated the efficacy of TAK-071 in rats with defined, partial losses of the cholinergic system, modeling the limited cholinergic losses seen in the patient groups listed above. Furthermore, loss of cholinergic neurons in rats has been frequently demonstrated to result in a loss of correct responses in cued, but not non-cued, trials of a sustained attention task (McGaughy et al., 1996; McGaughy and Sarter, 1998; Kucinski et al., 2013), consistent with the demonstration that fast cholinergic signaling is required for the detection of cues in such a task (Gritton et al., 2016; Sarter et al., 2016b; Howe et al., 2017). AChEIs do not benefit the performance of deafferented rats in this task (Kucinski et al., 2017). The present results indicate that administration of TAK-071 improves SAT performance exclusively in interaction with heightened demands on attentional control and, relatively more robustly in (female) rats exhibiting a propensity for reporting the presence of cues, that is, a relatively liberal response bias.

## MATERIALS AND METHODS

### Subjects

Adult male and female Sprague Dawley rats (Envigo, Indianapolis, IN), aged between 2 and 3 months of age at the binning of SAT training, were individually housed in opaque single standard cages (27.70 cm × 20.30 cm) in a temperature- and humidity-controlled environment (23°C, 45%). The final number of rats which reached SAT performance criterion, underwent successful surgery and reached the different stages of drug testing are indicated in Results.

The animals were maintained under a 12:12 hour light/dark schedule (lights on at 8:00 AM). Food (Rodent Chow; Envigo Teklad) was available *ad libitum*. Water access was gradually restricted over a 7-day period (12, 8, 5, 3, 1, 0.5, 0.25 hrs of water access per day) in the week before pre- and post-surgery behavioral testing. During testing, water was provided as rewards for correct responses during performance of the Sustained Attention Task (SAT). Rats were also provided water *ad libitum* for 20 min following each daily SAT session.

#### Body weights throughout the experimental period

Male rats were generally heavier than female rats, and male rats significantly increased their body weight throughout the experimental period. The analysis of the rats’ body weights prior to the onset of pre-surgery task training and the final week of drug testing (“time”) indicated main effects of sex (F(1,21)=124.71,*P*<0.001; males: 361.88±7.80 g; females: 267.31±3.84 g) and time (F(1,21)=7.88,P=0.01; pre-SAT: 305.60±9.59 g; final week of dosing: 319.80±12.06 g). A significant interaction between the effects of time and sex (F(1,21)=6.12,*P*=0.02) reflected that males, but not females, added weight during the testing period (males pre-SAT: 348.00±9.35 g; final week: 375.75±9.27 g; t(11)=7.45, *P*=0.02; females pre-SAT: 266.46±3.67 g, final week: 268.15±4.84 g; t(12)=0.19, *P*=0.67). Importantly, this sex-specific effect on body weights did not differ between sham-lesioned and lesioned rats (main effect of group: F(1,21)=1.78, *P*=0.20; 2- and 3-way interactions between group, sex, and time: all F<1.30, all *P*>0.26).

### Overall timeline of experiments

Animals were first trained on the SAT for between 2 and 5 months, depending on individual performances, prior to receiving lesions. Upon reaching the final stage of training, animals were required to reach criterion performance (detailed below) for 5 consecutive days before receiving cholinergic or sham lesions. We previously demonstrated that rats’ circadian rhythm entrained to daily SAT practice, yielding a robust diurnal phenotype (Gritton et al., 2009; Paolone et al., 2012; Gritton et al., 2013). Therefore, all behavioral testing likely occurred during the animals’ active phase of the day. Following training, animals underwent stereotaxic lesion surgeries followed by 2 weeks of recovery. During the final week of recovery animals were gradually water restricted. Post-surgery asymptotic SAT performance was determined over 10 days/sessions. Thereafter, drug effects (TAK-071; vehicle, 0.1 mg/kg, and 0.3 mg/kg) were assessed. Each dose was administered for 6 days during which regular SAT performance, the performance in the presence of a distractor (dSAT) and following the termination of the distractor (post-dSAT) were recorded (details below).

### Sustained Attention Task (SAT)

#### Apparatus

Training and testing were conducted using 12 operant chambers (MED Associates Inc.) housed within individual sound-attenuating cubicles. Throughout the duration of the experiments males and females were tested in separate chambers. Each chamber was equipped with two retractable levers, a central panel white light (2.8 W), and a water dispenser located on the same wall as the panel lights. The water dispenser delivered 45 µL of water per corrrect response. Signal presentation, lever operation, reinforcement delivery, and data collection were controlled by a Pentium PC and Med-PC for Windows software (version 4.1.3; MED Associates).

#### Acquisition

Water-deprived rats were initially trained to press a lever for a water reward in accordance with a modified fixed ratio-1 (FR1) schedule for water reinforcement. During this phase of training, any lever press resulted in the delivery of water. Typically, the animals do not exhibit a side bias with regard to which lever is pressed; however, if one lever was pressed 5 times in succession, the FR1 schedule was modified to require the animal to press the opposite lever before the next reward could be obtained. After 3 consecutive days with 120 reinforced lever presses each, the rats began training to discriminate between a signal (1 second illumination of the central panel light) and a non-signal (no illumination) event. Two seconds after a signal or non-signal event, both levers were extended into the operant chamber and remain extended for 4 s or until a lever was pressed. If no press occurred after 4 s, the levers retracted and an omission was recorded. Immediately following responses (either correct or incorrect), both levers were retracted and the variable intertrial interval (ITI; 12±3 s) was reset. On signal trials, a press of the left lever was reinforced and termed a “hit,” whereas a press of the right lever was not reinforced and termed a “miss.” On non-signal trials, a press of the right lever was reinforced and termed a “correct rejection,” whereas a press of the left lever was not reinforced and termed a “false alarm.” Animals received water rewards only for correct responses (45 µL for each hit and correct rejection), whereas incorrect responses (misses and false alarms) were not rewarded. To eliminate the possibility of a selection bias, half of the animals were trained with the opposite pattern. Signal and non-signal events were presented in pseudo-random order for 81 trials each (total of 162 trials) per session. During this phase of training, incorrect responses were followed by correction trials in which the previous trial was repeated. After three consecutive incorrect responses on correction trials, the animal underwent a forced trial in which the lever was extended for 90 s or until the animal made a response. If the forced choice trial was a signal trial, the signal light remained illuminated for as long as the lever was extended. The house light was not illuminated during this training stage.

Animals progressed to the subsequent step of shaping if they responded correctly to ≥70% of both signal and non-signal trials for 3 consecutive days. During the third phase of shaping, multiple signal durations (500, 50, and 25 ms) were introduced and the ITI was reduced to 9±3 s. Correction and forced-choice trials were also eliminated. Trial type and signal duration were pseudo-randomly determined for each trial. Session length was set at 40 min. After at least 3 days of stable performance, defined by at least 70% hits to 500 ms signals, 70% correct rejections, and ≤30% omissions, animals began training in the final version of the task. The final version was identical to the previous training stage except that the house light was illuminated throughout the session. The addition of the illuminated house light represents a crucial element of testing sustained attention as it requires the animal to constrain its behavior and focus on the central panel light during task performance. Upon reaching the final stage of training prior to lesion surgeries, animals remained at this stage until they reached a stable performance of at least 70% hits to 500 ms signals, 70% correct rejections, and ≤30% omissions for five consecutive sessions. Scores from the final 3 of these days were averaged to determine pre-surgery scores for each animal.

Animals practiced the dSAT a minimum of two times prior to surgery. In the dSAT, the first 8-min block of trials was identical to the SAT described above. This block was followed by a 16-min block with the distractor (chamber houselight flashing on/off at 0.5 Hz) turned on (dSAT). This distractor profoundly disrupts performance, yielding chance level performance. After distractor termination, post-dSAT performance was determined during a final 16-min block of the regular SAT. Individual distractor test sessions were separated by a minimum of 5 days of regular SAT practice sessions. Our collective evidence indicates that repeated exposure to this distractor does not significantly reduce the efficacy of the distractor, including the rate of post-distractor performance recovery.

#### Post-surgery performance and drug administration regimen

Two weeks following surgeries animals were water-deprived over 1 week in the same manner as described above and then tested for 10 consecutive days on the SAT to determine the effects of cholinergic or sham lesions on performances. A dSAT session was given on the 6^th^ day. SAT performance on the 8^th^, 9^th^ and 10^th^ days was averaged to determine post lesion scores. Thereafter, drug effects were tested in blocks of 6-days per dose. The order of dosing was pseudo-randomized to control for potential sequence effects and individual 6-day blocks were separated by one week of daily SAT practice without vehicle or drug administration. From each 6-day block, data from regular SAT sessions on the 3^rd^, 4^th^ and 5^th^ days were averaged to determine drug or vehicle effects. dSAT performance was determined on the 6^th^ day.

#### Measures of SAT performance

The following behavior measures were recorded during each SAT session: hits, misses, false alarms, correct rejections, and omissions. Misses and false alarms are the inverse of hits and correct rejections, respectively. The relative number of hits (hits/hits+misses) for each signal length as well as the relative number of correct rejections (correct rejections/correct rejections+false alarms) were calculated. In addition, an overall measure of attentional aptitude, the SAT score, that integrates both the relative number of hits (h) and the relative number of false alarms (f), was also determined at each signal duration. The SAT score was calculated using the following formula: (h-f)/[2(h+f)-(h+f)^2^] (Frey and Colliver, 1973).

Thus, SAT scores are not confounded by errors of omission, which were analyzed separately. SAT scores range from 1.0 to - 1.0, with 1.0 indicating that all responses were hits and correct rejections, 0 indicating an inability to discriminate between signal and non-signal events, and −1.0 indicating that all responses were misses and false alarms. Furthermore, SAT scores ≤0.17 indicate chance performance (for signal and non-signal trials, and assuming equal probability for the two trial outcomes, 59/100 successes yield a probability of just under *P*=0.05, and a SAT score of 0.18). Scores between 0.18 and 0.51 (above chance but less than 3 fold higher than chance) are considered to indicate “medium level” of SAT performance, while scores above 0.51 (over 3 fold higher than chance), typically seen only for SAT scores calculated over hits to longest signals, indicate a “high level” of SAT performance.

#### Signal detection parameters

To further determine the nature of the effects of TAK-071 on post-dSAT performance in sham-lesioned and lesioned male and female rats, we determined perceptual sensitivity (*d’*) and bias (*B”*_*D*_) based on the proportion of hits (H_P_) and proportion of false alarms (FA_P_). Combining hits across signal duration, *d’* was calculated using the formula *d’*=z(H_P_)–z(FA_P_) (Green and Swets, 1974). High values of *d’*, maximum of 4.65, occur when all signals are correctly reported and very few false alarms occur. *d’* is zero when the proportion of hits is equal to the proportion of false alarms. *B”*_*D*_ was similarly calculated across signal durations using the formula: *B”*_*D*_=[(1-H_P_)(1-FA_P_) -H_P_FA_P_]/[(1-H_P_)(1-FA_P_) +H_P_FA_P_] (Donaldson, 1992). *B”*_*D*_ ranges from −1 to +1 with negative values indicating a liberal bias towards reporting the presentation of a signal and positive values indicating a conservative bias. A *B”*_*D*_ measure of 0 indicates no bias (for a prior analysis of these parameters based on data from humans performing the SAT and dSAT see Demeter et al., 2008).

### Lesions

Cholino-specific basal forebrain lesions were carried out with the goal to generate a 40-60% decrease of the density of the cortical cholinergic inputs system. Such relatively moderate losses were expected to yield decreases in hit rates without abolishing signal duration-dependent performance, preserving a degree of performance-associated ACh release, and thus preserving post-synaptic muscarinic receptor function (Rossner et al., 1995a; Rossner et al., 1995b; Fadel et al., 1996; McGaughy et al., 1996; McGaughy and Sarter, 1998; Dalley et al., 2001; Kucinski et al., 2013; Kucinski et al., 2017). Rats with such moderate cholinergic losses may model the impairments in attention in patients with such limited cholinergic losses and whose cognitive capacity overall remain comparable with healthy controls (Kim et al., 2017b).

Rats were first placed in vaporization chambers and anesthetized with 4-5% isoflurane (delivered at 0.6 L/min O2; SurgiVet Isotec 4 Anesthesia Vaporizer) until the animals were no longer responsive to a tail pinch and exhibited no hind-limb withdrawal reflex. The animals’ heads were shaved using electric clippers and cleaned with a betadine scrub. The animals were then mounted to a stereotaxic instrument (David Kopf Instruments) and isoflurane anesthesia was maintained at 1-3% for the remainder of the surgery. The animals’ body temperature was maintained at 37°C using Deltaphase isothermal pads (Braintree Scientific). Ophthalmic ointment was provided for lubrication of the eyes. To prevent hypovolemia and hemodynamic instability during prolonged surgeries, 1 mL/100 g 0.9% of NaCl (s.c.) was administered. Animals also received an injection of an analgesic (Carprofen; 5.0 mg/kg; s.c) prior to surgery and once or twice daily as necessary for 48 hours post-operatively. Basal forebrain cholinergic neurons situated in the nucleus basalis and substantia innominata were targeted with the immunotoxin 192 IgG-saporin (SAP; Advanced Targeting Systems) in aCSF (Tocris Bioscience, Bristol, United Kingdom) infused bilaterally (120 ng/µL; 0.5 µL/hemisphere; AP −0.8; ML ± 2.9; DV −7.8). Sham-surgeries were conducted by infusing equal volumes of artificial cerebrospinal fluid. Following infusions the needle was left in position for 5-10 min to foster absorption of the toxin. Non-absorbable sutures were used to close the incisions and a topical antibiotic (Neosporin) was applied to the wounds immediately after surgery.

### TAK-071

TAK-071 (Takeda Pharmaceutical Company; Lot # M071-004) (Sako et al., 2018; Kurimoto et al., 2019) was suspended in 0.5% methyl cellulose in DI water (vehicle). Vehicle and a 0.1 mg/mL stock solution for each dose of TAK-071 were prepared fresh every week. Drug or vehicle were administered by oral gavage. The volume of drug or vehicle administered to male and female rats ranged from 0.4-1.2 mL and 0.3-0.9 mL, respectively (see *Subjects* for body weights).

Animals were extensively familiarized with gavage administration procedures during the post-surgery recovery period and the period during which asymptotic post-surgery SAT performance was determined. TAK-071 was administered at 0.1 and 0.3 mg/kg (see *Post-surgery performance and drug administration regimen*, above, for details of the administration regimen). A small subgroup of rats were also administered smaller doses of TAK-071 (0.01 and 0.03 mg/kg) but these data were excluded from the final analyses. Drug or vehicle was administered 2 hours prior to SAT sessions (e.g., administration at 9 a.m., followed be SAT testing 11:00-11:45 a.m.).

Pharmacokinetic data were previously reported (see Supplementary Information in Sako et al., 2018). These data indicate that following the oral administration of 0.1 mg/kg TAK-071, plasma levels, at the time of SAT testing in the present experiment (120-165 min post administration), remained near the maximum concentration (Cmax: 85 ng/mL; Tmax: 1.9 hrs; see Table S2c and Fig. S4e in Sako et al. 2018).

### Histological methods and quantification of cortical AChE-positive fiber density

Following the completion of post-surgery drug testing, rats were deeply anesthetized and transcardially perfused at a rate of 50 mL/min with 0.1M phosphate buffer solution (PBS) for 2 min followed by perfusion with 4% paraformaldehyde in 0.4M sodium-phosphate solution and 15% picric acid (pH 7.4) for 9 min. Brains were rapidly removed and postfixed for 2-6 h at 4°C and then rinsed in 0.1M PBS and stored in 30% sucrose solution and allowed to sink. Coronal sections (40 µM thickness) were sliced using a freezing microtome (CM 2000R; Leica) and stored in antifreeze solution.

To visualize acetylcholine esterase-(AChE) positive fibers, sections were first rinsed in 0.1M PBS three times for 15 min each before a 30 min incubation in 0.1% hydrogen peroxide-0.1M phosphate buffer solution. Sections were then rinsed 3 × 5 min with maleate buffer (3.14g NaOH, 5.83g maleic acid, per 1L diH20, pH 6.0). Next, sections were incubated for 75 min in a solution of maleate buffer (200 mL), 0.1M sodium citrate (0.5 mL), 30 mM cupric sulfate (1.0 mL), 5 mM potassium ferricyanide (1.0 mL) and 0.5 mg of acetylthiocholine iodide (0.5 mg per section). Sections were then rinsed 3 × 5 min with 30mM TRIS solution (3.12g trizma base, 200 mL of 0.1N HCl, in 1L diH20, pH 7.60). Sections were then incubated in a solution of 30mM TRIS (250 mL), ammonium nickel sulfate (0.75g), and DAB (3,3’-diaminobenzidine) (100 mg), pH 6.30, for 10 min. Next, 8.0 uL of 30% H_2_O_2_ per 40 mL of solution was added and sections were incubated until they became dark black (approx.. 90 s). Sections were then rinsed in 3 mM TRIS solution 3 × 5 min before being mounted on gelatin-coated slides and allowed to dry overnight. The following day, slides were dehydrated in an ascending alcohol series (70%, 90%, and 100% ethanol) and defatted in xylene before cover-slipping.

Photographs of the AChE stained sections were taken at 40x magnification using a Leica DM400B digital microscope, Leica DMC6200 camera, and Leica LAS X software. For each rat, one section at approximately Bregma AP −0.8 mm was used to generate an estimate of lesion effects by assessing AChE-positive fiber density in layers V and VI of the primary motor cortex. Fibers were counted in both hemispheres. A 4 × 4 square grid, covering an area of 150 um^2^, was imposed on each of the photographs. AChE-positive fibers were counted at each point where they crossed one of the lines of the grid (see Figure 4 for illustration of this method; see also McGaughy et al., 1996; Burk and Sarter, 2001).

### Statistical analyses

SAT and dSAT performances were analyzed primarily by using mixed-model repeated-measures ANOVAs as well as one or two-way ANOVAs when applicable. Sex was a factor in all analyses. The analysis of SAT scores and hits also included the within-subject factor signal duration (500, 50, and 25 ms). Following significant main effects, post hoc multiple comparisons were conducted using the Least Significant Difference (LSD) test. Significant interactions between the effects of group and other factors were followed by one-way ANOVAs on the effects of group or t-tests and LSD multiple comparison tests. Statistical analyses were performed using SPSS for Windows (version 17.0: SPSS). In cases of violation of the sphericity assumption, Huyhn–Feldt-corrected F-values, along with uncorrected degrees of freedom, are given. Alpha was set at 0.05. For the analysis of effects of lesions and drug in SAT, dSAT and post-dSAT performance, the SAT score was the primary outcome measure. Secondary analyses concerned effects on hits and correct rejections to locate, *post hoc*, the main source(s) of effects on the SAT score. Therefore, the possibility of cumulative type I errors, arising from the analysis of multiple dependent measures (SAT score, hits, correct rejections), was not taken into account. For data analyzed using parametric tests, variances were reported and illustrated as standard error of the mean (SEM). To determine the direction and strength of potential monotonic relationships between performance parameters and AChE-positive fiber counts Spearman’s *rho* was determined. Non-parametric statistical test (Mann-Whitney U test, Wilcoxon’s matched pairs test) were used to determine lesion effects on AChE-positive fiber counts and effects of lesions and the treatment on the joint probabilities for hits in trials that were part of certain trial sequences (see Results). Exact *P* values are reported as recommended previously (Greenwald et al., 1996; Sarter and Fritschy, 2008).

## RESULTS

### Pre-surgery performance

Prior to surgeries, rats (N=25, 13 females) underwent SAT training until they reached stable criterion performance. Rats required on average 13.89±1.13 days to complete the first stage of training (range 7 to 28 days), 10.24±1.55 days to complete the second stage (range 3 to 35 days), and a total of 47.96±5.05 days to reach SAT criterion performance (range 15 to 94 days). The number of days required to complete the 3 stages of training did not differ between groups (assigned to sham versus SAP-lesion surgery; n=12 and 13, respectively) or sex (12 males, 13 females; main effects and interactions: all F<1.66, all *P*>0.21).

Performance data from the final 3 pre-surgery SAT sessions were averaged. Overall, as expected, there was a significant main effect of signal duration on the SAT score and hit rates (both F(2,42)>208.26, both *P*<0.001; Fig. 1a,b). There were no effects of sex and group (forthcoming sham or SAP surgery) on any of the measures of performance (main effects and interactions: al; F<2.98, all *P*>0.10). Thus, prior to surgery, SAT performance did not differ between rats assigned to sham versus SAP lesions performance.

**Figure 1.**
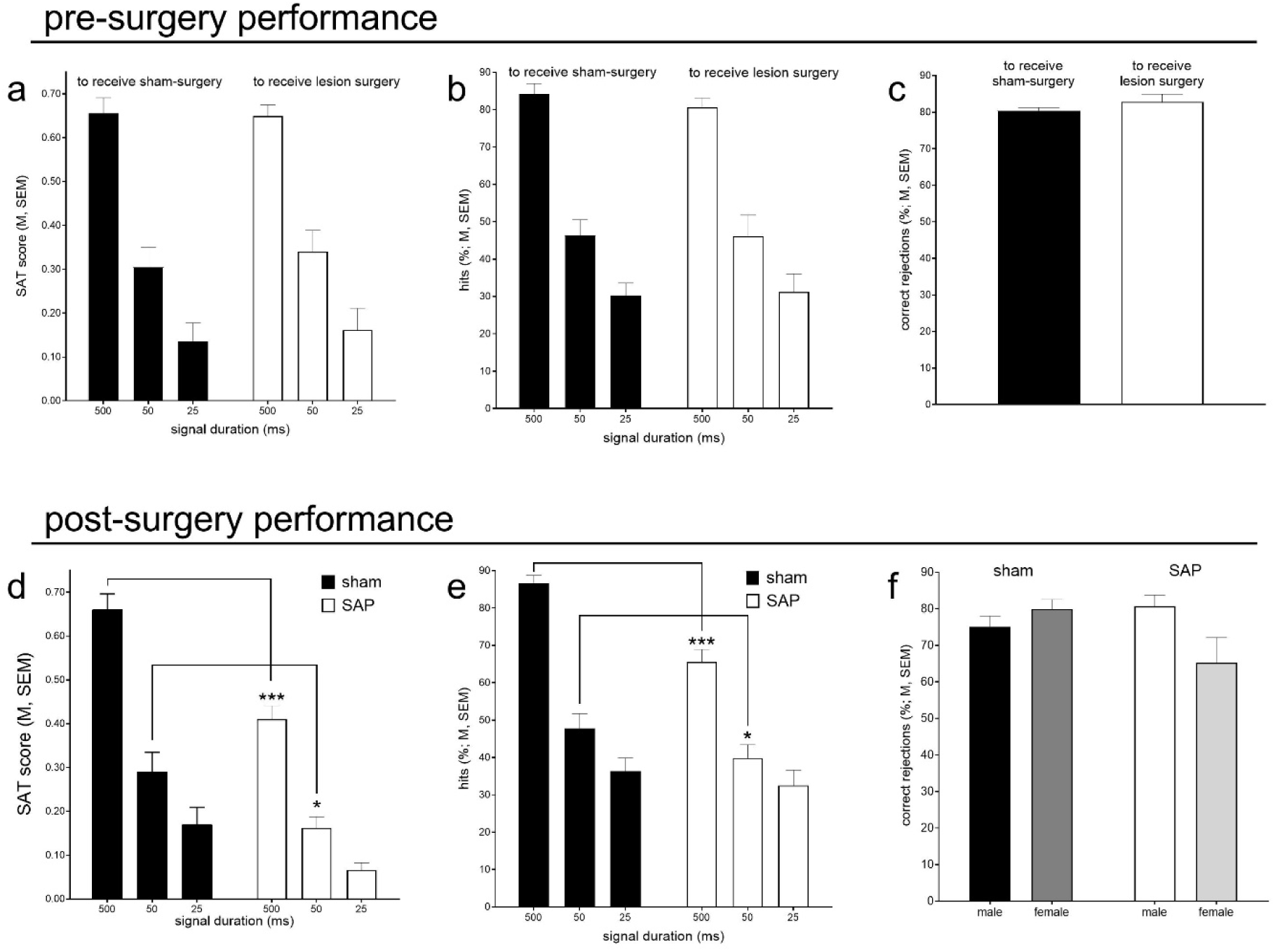
Prior to surgeries, the SAT performance of rats assigned to sham versus SAP lesion surgery did not differ statistically. As expected, SAT scores and hits varied by signal duration (main effects), indicating that SAT scores (<u>a</u>) calculated on the basis of hits to 500 ms signals were significantly higher than scores computed over hits to 50 and 25 ms signals, and SAT scores calculated on the basis of 50 ms signals were higher than those over 25 ms signals (*post hoc* comparisons: all P<0.001). Likewise, hit rates (<u>b</u>) differed between all three signal durations (all *P*<0.001). Rats omitted less than 4% of all trials. There were no effects of sex on pre-surgery performance (data not shown). Lesions of the cholinergic system impaired SAT performance (<u>d</u>) which reflected a selective decrease in hit rates (<u>e</u>; see Results for ANOVAs; for this and all subsequent figures, *post hoc* multiple comparisons are indicated: *,**,***, *P*<0.05, 0.01, 0.001). The lesions did not affect the correct rejection rate. A significant interaction between the effects of group and sex on correct rejections (f) may have indicated a lesion-induced adoption of a “riskier” criterion for reporting the presence of a cue in females (*post hoc* comparisons failed to locate this interaction).

### Effects of cholinergic loss on SAT performance

Following the completion of the post-surgery recovery period of 3 weeks, animals underwent 10 consecutive days of SAT testing to determine SAT performance following sham and SAP lesions. For this analysis, performance scores obtained on the 8^th^-10^th^ day were averaged for each individual rat.

The loss of cholinergic neurons impaired hit rates, primarily to longer signals, consistent with evidence from prior experiments on the effects of such lesions, and with the essential role of the cholinergic system in the detection of cues (see Introduction). The cholinergic lesions reduced SAT scores (main effect: F(1,21)=13.04, *P*=0.002; shams: 0.35±0.03; SAP: 0.21±0.02; Fig. 1d). Notably, the average SAT score for lesioned rats was close to chance performance (0.17; see Methods for the interpretation of this score). This effect of the lesions on the SAT score interacted significantly with the effects of signal duration (main effect: F(2,42)=251.34, *P*<0.001; interaction: F(2,42)=8.31, *P*=0.001), reflecting that scores calculated over hits to the two longer signal durations were significantly lower in lesioned when compared with sham-lesioned rats (Fig. 1d; results of *post hoc* multiple comparisons are indicated in the Figures).

As illustrated in Fig. 1e, the effects of the lesions on SAT scores were largely reproduced by effects on hit rates (main effects of signal duration, group and interaction: all F>6.91, all *P*<0.02). Moreover, however, females scored more hits than males (F(1,21)=6.89, *P*=0.02), irrespective of lesion status (group x sex: F(1,21)=1.27, *P*=0.27) and signal duration (2- and 3-way interactions: all F<0.07, all *P*>0.94; males: 45.47±2.93%; females: 55.08± 2.30%).

The about 10% greater hit rate in females was mirrored by about a 10% lower correct rejection rate in lesioned female rats (group x sex; F(1,21)=4.95, *P*=0.04; Fig. 1f; no main effects of group or sex, both F(1,21)<1.37, both *P*>0.26). *Post hoc* multiple comparisons failed to confirm that this interaction reflected a significant decrease in the correct rejection rates in lesioned females compared to lesioned males (shams: t(11)=1.80, P=0.21; lesioned: t(12)=3.50, *P*=0.09; Fig. 1f). The lesions had no effect on the relatively low number of errors of omission (<4% of trials; main effects and group x sex: all F(1,21)<0.74, all *P*>0.40). The relatively high hit and lower correct rejection rates in lesioned females suggested a relatively greater propensity to report the presence of signals, that is, a more liberal response bias (below).

### TAK-071 did not affect regular (unchallenged) SAT performance

At the beginning of the assessment of TAK-071 (see Methods for details of the administration regimen), one sham-lesioned rat completely stopped performing and a second sham-lesioned rat failed to be successfully familiarized with the administration procedure by oral gavage. Thus, drug effects on SAT, dSAT and post-dSAT are based on data from 10 sham-lesioned (5 females) and 13 lesioned (7 females) rats.

Administration of TAK-071 (0.1 or 0.3 mg/kg) did not affect unchallenged, regular SAT performance (all main effects and interactions with group, and sex and signal duration (where applicable): all F<2.30, all *P*>0.11).

The analysis of the effects of TAK-071 on regular (unchallenged) SAT performance reproduced the effects of the lesions and interactions with signal duration on SAT scores obtained from the post-surgery period (main effects of group and signal duration, signal × group: all F>3.90, all *P*<0.03). Likewise, the analysis of hits reproduced the group × signal duration interaction seen before (F(2,38)=9.45, *P*<0.001), but the main effect of group on hits did not reach significance (F(1,19)=3.15, *P*=0.09), reflecting an increase in the variability of the hit rates observed in lesioned rats when compared with the performance prior to administering drug or vehicle (above).

Overall, females performed better than males (SAT score: F(1,19)=1.27, *P*= 0.03; males: 0.30±0.02; females: 0.35±0.04), reflecting in part a trend for more hits in females (F(1,109)=4.25, *P*=0.053; males 47.73±4.23%; females: 58.50±3.04%). Moreover, the group x sex interaction on correct rejections, seen during the post-surgery period, was reproduced in this analysis (F(1,19)=5.34, *P*=0.03) and indicated that lesioned (t(12)=4.87, P=0.04), but not sham-lesioned, females (t(9)=1.28, P=0.29) scored less correct rejections (or, conversely, more false alarms) than their male counterparts. Once again, this evidence indicated a relatively greater propensity of lesioned female rats to report the presence of a cue.

### Performance disruption in the presence of the distractor (dSAT)

In the SAT version used for rats, the flashing houselight distractor acts as a perceptual disruptor rather than a psychological distractor (Sarter et al., 2016a). Furthermore, this disruptor is highly effective, yielding near or at chance performance in all rats. Accordingly, we did not expect the treatment with TAK-071 - or any pro-cholinergic treatment - to significantly attenuate the acute effects of this distractor (main effects of drug dose and interactions involving dose, on all measures of performance; all F<2.08, all *P*>0.14). Disruption of SAT performance serves primarily as a means to investigate the capacity for post-distractor performance recovery (below), as such recovery is thought to reveal attentional control capacities (Sarter and Paolone, 2011; St Peters et al., 2011; Berry et al., 2015; Kim et al., 2017b).

Confirming the ability of the distractor to robustly depress all rats’ performance, the effects of the lesions and the interactions between group and signal duration on SAT scores and hit rates, which were seen in the absence of the distractor, were abolished during dSAT (main effects of group and interactions with signal duration: all F<3.11, all *P*>0.09; Figure 2a). Indeed, SAT scores, with the exception of those calculated over hits to longest signals in sham-lesioned rats, indicated chance performance (Fig. 2a). In the presence of the distractor, main effects of signal duration on the SAT score and on hit rate were preserved (both F>23.18, both P<0.001), reflecting that performance in the presence of longest signals continued to be more accurate than responding to 50 and 25 ms signals (*post hoc* multiple comparisons: all *P*<0.001) but responding to the 50 ms signal no longer was more accurate than to the 25 ms signal (both *post hoc P*<0.08). Omissions remained below 6% of all trials and for all animals (all *P*>0.21).

**Figure 2.**
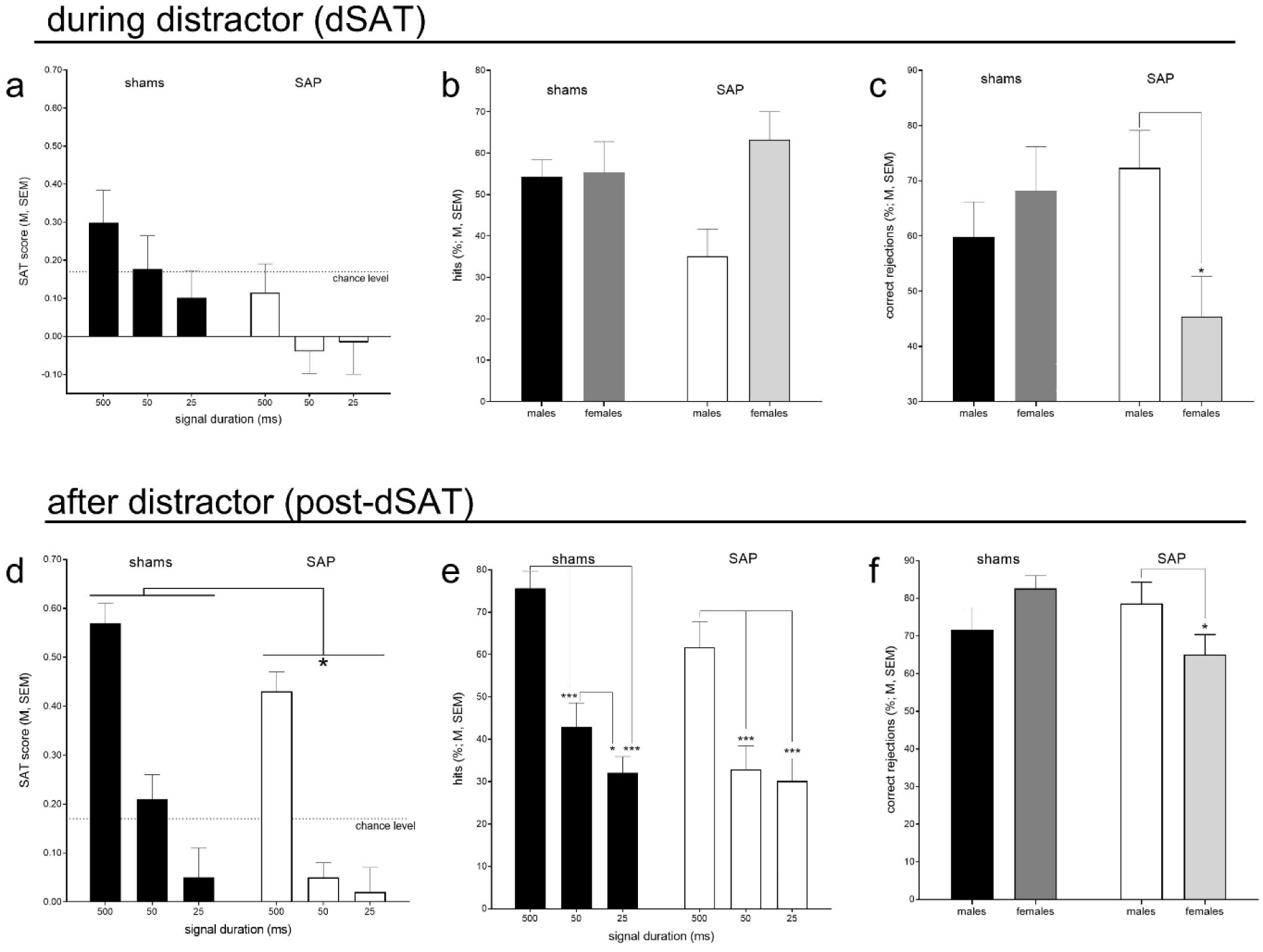
As was expected, the distractor reduced performance of all rats to chance levels of performance (see Methods for definition; see dotted line in <u>a</u>). Accordingly, the effects of lesions, seen in the absence of the distractor, were no longer statistically apparent during dSAT. In the analysis of SAT scores and hits, signal duration-dependent performance was preserved during dSAT. TAK-071 neither was expected nor found to affect dSAT performance. In the presence of the distractor, females scored more hits (main effect) and, as illustrated in <u>b</u>, this reflected a trend (*P*=0.054) for lesioned males to miss a relatively large proportion of hits when compared with lesioned females. Furthermore, a significant interaction between the effects of sex and group on correct rejections (<u>c</u>) reflected that lesioned females responded to the distractor by scoring relatively less correct rejections than males (or, conversely, more false alarms). In conjunction with their higher hit rates, lesioned females adopted a relatively a more liberal response bias than lesioned males during dSAT. During the post-distractor period, the performance of sham rats recovered more robustly than that of SAP rats, as indicated by the re-emergence of a group effect on the SAT (<u>d</u>; data in d are shown by signal duration to foster visual comparisons with SAT scores during dSAT (<u>a</u>) and pre- and post-surgery (Fig. 1a,d)) and a significant interaction between the effects of group x signal duration on hits (<u>e</u>; multiple comparisons failed to locate the interaction and indicated a trend (P=0.09) for more hits to 500 ms signals in sham rats). During the post-dSAT period, lesioned females again scored less correct rejections (or me false alarms than females (f) although the effect appeared less pronounced than in the presence of the distractor (<u>c</u>).

In the presence of the distractor, the manifestation of opposite response biases of males and female rats appeared even greater than during regular SAT performance, as indicated by a main effect of sex on hits (F1,19)=4.96, *P*=0.040) and reflecting a near 17% higher hit rate in females (males: 43.82±4.90; females: 60.00±4.90). Moreover, a trend for a sex × group interaction on hits (F(1,19)=4.24, *P*=0.054; Fig. 2b) appeared to reflect that lesioned males scored less hits than sham males in the presence of the distractor, while lesioned females tended to increase their hit rate relative to their sham-lesioned counterparts.

Concerning correct rejections during dSAT, the group × sex interaction again was significant (F(1,19)=5.85, *P*=0.03; Fig. 2c). Compared with the animals’ correct rejection rates during regular SAT, during dSAT, this interaction appeared to be based on a relatively larger asymmetry between the correct rejection rates of lesioned males and females. This observation was confirmed by a *post hoc* analysis of the effects of sex and task type (SAT versus dSAT) on correct rejections in sham and SAP-lesioned rats. In sham-lesioned rats, a main effect of the distractor (F(1,8)=19.52, *P*=0.002; SAT: 78.20±2.78%; dSAT: 64.00±5.02%) did not interact with sex (main effect of sex and interaction, both F<0.91, both *P*>0.37). In contrast, in lesioned rats, significant effects of task type and sex, and a significant interaction between the two factors (all F(1,11)>6.29, all P<0.03) reflected that, in the presence of the distractor, the correct rejection rate of lesioned females decreased more robustly than in males. Together with the higher hit rates in females during dSAT performance, these *post-hoc* analyses suggested that female rats, and particularly lesioned females, adopted a more liberal response criterion (more hits, but also more false alarms) than males when performance was taxed by the distractor.

### TAK-071 improves post-dSAT performance

Following the termination of the distractor, performance of sham-lesioned rats during the subsequent 16-min post-dSAT period recovered to a greater degree than in SAP rats, as indicated by a main effect of group on the SAT score (F(1,19)=4.60, *P*=0.045; shams: 0.28±0.05; SAP: 0.16±0.03; note that the average SAT score of lesioned rats indicated chance performance; Fig. 2d). Post-dSAT performance recovery was previously demonstrated to depend on cholinergic activity (St Peters et al., 2011; Berry et al., 2015; Kim et al., 2017b). A signal duration x group interaction on hits (F(2,38)=4.13, *P*=0.02; Fig. 2e) further suggested that, in sham rats, hits to longer signals recovered more greatly than hits to shorter signals (*post hoc* multiple comparisons only indicated a trend for more hits to longest signals in shams versus SAP rats; t(22)=3.21, *P*=0.09). Once again, lesioned females scored less correct rejections than lesioned males (group x sex: F(1,19)=5.27, *P*=0.03; Fig. 2e) although, in the absence of the distractor challenge, this effect appeared to be less pronounced than during dSAT (Fig. 2c).

Treatment with TAK-071 improved the post-dSAT performance of all rats. A main effect of dose on SAT (F2,28)=5.48, *P*=0.008) was based primarily on an increase in performance following the lower dose of the drug (Fig. 3a; no interaction between dose and signal duration: F(4,76)=0.51, *P*=0.73). Administration of 0.1 mg/kg of TAK-071 yielded a medium-to-large improvement in performance (Cohen’s *d*=0.64).

**Figure 3.**
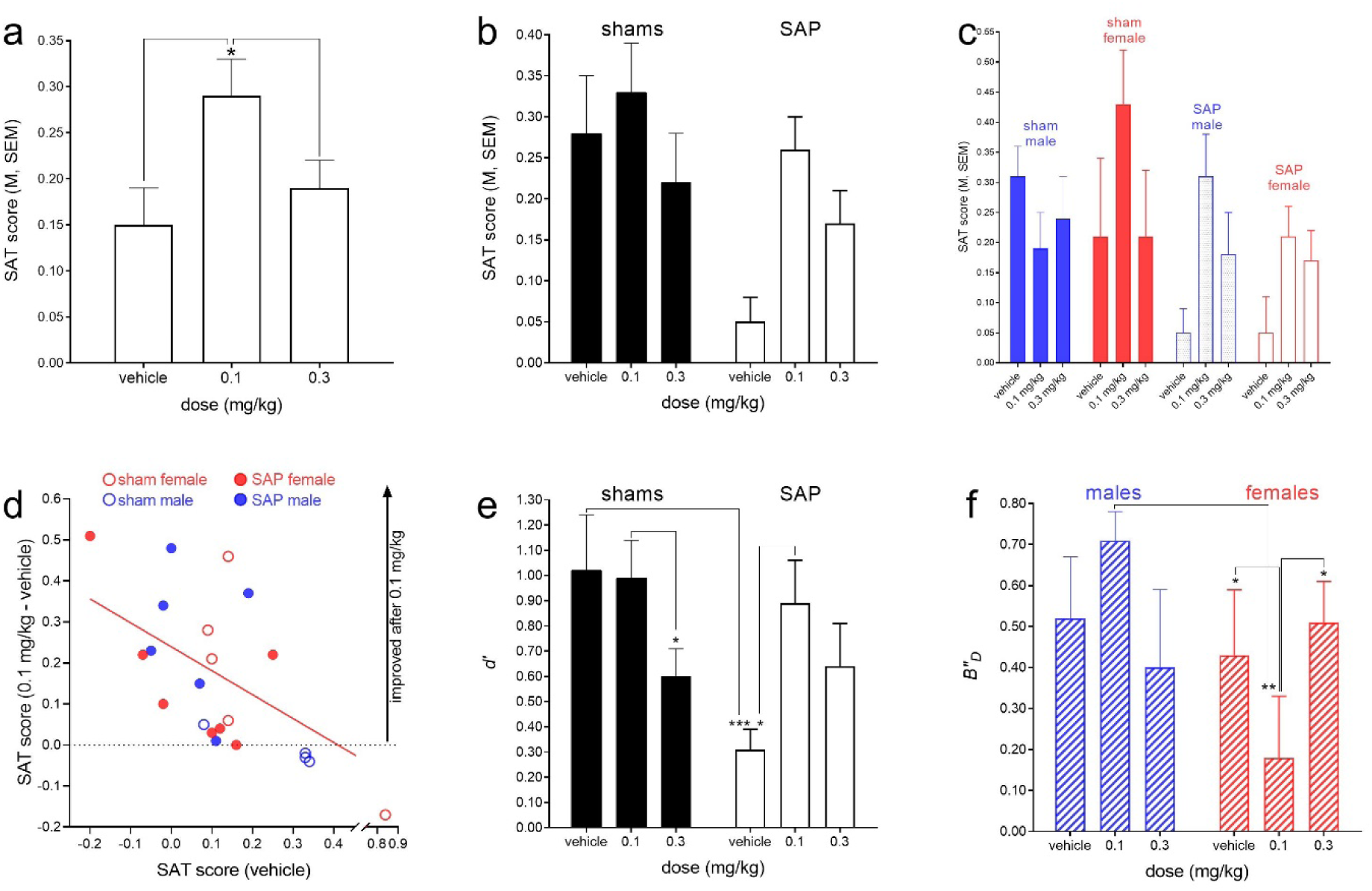
Administration of TAK-071 improved the performance of all rats during the 16-min post-distractor performance period (post-dSAT; <u>a</u>; see Results for effect size). Additional analysis indicated a trend for a relatively greater effect of the drug in lesioned over sham-lesioned rats (<u>b</u>), and that both doses improved the performance of lesioned rats. Moreover, in lesioned rats, both doses improved the performance of both males and females, while the lower dose also appeared to benefit the SAT scores of sham-lesioned females (<u>c</u>), perhaps associated with these animals’ relatively more liberal response bias (see Discussion). Reflecting that drug effects were relatively more robust in lesioned rats, the efficacy of the lower dose was highly correlated with the SAT score in the absence of drug (<u>d</u>; note that removal of the outlier at the lower right did not affect this conclusion; see Results). Analysis of signal detection theory-derived measures of perceptual sensitivity *d’* (<u>e</u>) and response bias *B”*_*D*_(f) indicated that the lower dose of the drug enhanced the sensitivity of all lesioned rats but, in addition, benefited the performance of female rats by fostering the adoption of a more liberal bias, thought to reflect an adaptive, top-down-orchestrated propensity to report the presence of cues.

The analysis of the effects of TAK-071 on SAT scores obtained during the post-dSAT period also indicated a trend for an interaction between the effects of dose and group (F(2,38)=5.48, *P*=0.06). This trend reflected that 0.1 mg/kg TAK-071 rescued the performance in lesioned rats more robustly than in control rats (Fig. 3b).

Moreover, inspection of SAT scores from sham and lesioned males and females indicated that TAK-071 increased dSAT scores in both lesioned males and females but in sham-lesioned rats, only in females (Fig. 3c). However, the analysis of the 3-way interaction did not reach significance (dose × group × sex; F(2,38)=2.75, *P*=0.08), due in part to the relatively high variability of SAT scores in sham-lesioned females. The data depicted in Fig. 3c further suggested that both the 0.1 and 0.3 mg/kg doses of drug increased SAT scores in lesioned rats while, in sham-lesioned females, only the smaller dose improved performance. A trend for an interaction between the effects of dose and sex on hit rates (F(2,38)=3.15, *P*=0.054; males, vehicle, 0.1, 0.3 mg/kg (means, averaged over all signal durations): 39.73%, 39.76%, 42.18%; females: 44.26%, 59.69%; 44.11%) further suggested that TAK-071 generally was more effective in female rats. TAK-071 did not significantly affect the relatively number of correct rejections and errors of omission (all main effect and interactions involving dose: F<2.47, all P>0.10).

Taken together, the analysis of the effects of TAK-071 indicated efficacy only during the post-dSAT period, robust efficacy of the 0.1 mg/kg dose in lesioned and non-lesioned rats (main effect), and strong trends indicating relatively greater efficacy of the drug in lesioned over sham-lesioned rats, and in female over male rats. These trends were also mirrored in the distribution of data that supported a highly significant correlation between the SAT scores from vehicle treated rats and the difference between scores obtained following the treatment with 0.1 mg/kg and vehicle scores (Spearman’s *rho*; ρ=-0.63, *P*=0.002; Fig. 3d; note that the correlation remains significant if the bottom-right outlier is removed, ρ=-0.58, *P*=0.006). This correlation indicated that, generally, lower SAT scores in the absence of drug predicted greater effects of the 0.1 mg/kg dose, consistent with greater effects seen in lesioned rats (see filled circles in Fig. 3d; also note efficacy in sham-lesioned females, open red circles in Fig. 3d).

#### Analysis of d’ and B”_D_ indicated multiple sources of drug effects

We determined, *post hoc*, the signal detection theory-derived parameters of perceptual sensitivity *d’* and bias *B”*_*D*_ (Green and Swets, 1974; Donaldson, 1992), to gain insight into the perceptual versus cognitive nature of the effects of TAK-071. The analysis of *d’* reproduced the main effect of dose on the SAT score (shown in Fig. 3a), consistent with the view that enhancing cholinergic function primarily acts to increase cue detection rates (F(2,38)=4.71, *P*=0.02; vehicle: 0.62±0.13; 0.1 mg/kg: 0.94±0.11; 0.3 mg/kg: 0.62±0.11; no main effects of group or sex; both F<2.12, both *P*>0.16). Moreover, the effects of dose and group interacted significantly (F(2,38)=6.15,*P*=0.005), reflecting that in shams, the higher dose of TAK-071 reduced perceptual sensitivity while in lesioned rats, the lower dose significantly enhanced it (Fig. 3e). However, drug effects on perceptual sensitivity were unrelated to the sex of the rats (effects of sex, and interactions involving sex and dose: all F<2.00, all P>0.15).

A reversed situation was indicated in the analysis of the response bias *B”*_*D*_. There were no effects of dose, group, sex, and no interactions between dose and group (all F<0.75, all *P*>0.40). However, a significant dose x sex interaction (F(2,38)=5.28,*P*=0.009) indicated that all females (dose x sex x group: F(2,38)=0.61,*P*=0.55) adopted a significantly more liberal bias following the administration of 0.1 mg/kg than following vehicle or the higher dose. In contrast, following the administration of the lower dose, males adopted a highly more conservative bias than females (Fig. 3f). Thus, females appeared to have benefited not only from a drug effect on perceptual sensitivity - an effect they shared with all lesioned rats - but also from a generally greater drug-induced propensity to report the presence of a cue.

#### TAK-071 increased shift-hits over consecutive hits

We previously demonstrated that cholinergic transients mediate the detection of cues, that is, hits in the SAT (Howe et al., 2013). Moreover, however, these transients only occurred during hits that followed blank trials (factual or perceived) and not during hits that followed cued trials (factual or perceived). This finding indicates that cholinergic transients specifically mediate cue detection if involving a shift from attentional monitoring to cue-oriented behavior (“shift-hits”), but that transients are not necessary when that shift already occurred (“consecutive hits) (Sarter and Lustig, 2019). This hypothesis predicts that cholinergic losses reduce shift-hits to a greater degree than consecutive hits, and that a drug which rescues attenuated cholinergic activity improves primarily shift-hits. To test these predictions, we determined *post-hoc*, and based on the post-dSAT data, the joint probabilities for a cued trial to follow a hit (h) or a false alarm (fa), and for this cued trial to yield a hit (*p(hit+fa,hit*)), and for a cued trial to follow a correct rejection (cr) or a miss (m), and for this cued trial to yield a hit (*p(cr+m,hit*). This analysis excluded the data from two SAP rats which did not allow the calculation of all probabilities because at least one divisor was zero (n=11 shams and 11 SAP rats). Rejecting the first prediction, the decrease in hits after cholinergic losses could not be attributed solely to a reduction in the joint probability for shift-hits (shams: *p*=0.26; SAP: *p*=0.21 (medians); Mann-Whitney U-test: U=33.50, *P*=0.078; consecutive hits: shams: *p*=0.24; SAP: *p*=0.17; U=55.00, *P*=0.73). However, in SAP rats, administration of 0.1 mg/kg TAK-071 increased the probability for shift-hits (vehicle: *p*=0.21; 0.1 mg/kg: *p*=0.26; Wilcoxon’s matched-pairs test: W=56.00, *P*=0.0098), but not for consecutive hits (*P*=0.64). TAK-071 neither affected the probabilities for shift-hits nor for consecutive hits in sham-lesioned controls (both *P*>0.70). These results are consistent with a pro-cholinergic mechanism of TAK-071 (see Discussion).

### AChE-positive fiber counts and relationships with performance and drug effects

Figure 4 illustrates the effects of infusions of 192 IgG-saporin into the basal forebrain on the density of cholinergic axons and varicosities in cortex and the grid counting method used to estimate the cholinergic losses in the cortex.

Fiber counts were obtained from the cortex of both hemispheres. Although the hemispheric counts were highly correlated (ρ=0.58, *P*=0.0047), high versus low counts differed significantly (t(42)=2.30, P=0.03; low counts: 56.55±5.82 counts; high counts: 76.45±6.42). We therefore analyzed low and high counts separately (Fig. 5), in part guided by the assumption that the low count more validly indicates the degree of impairment resulting from bilateral lesions. Both low and high counts differed significantly between sham and SAP-lesioned rats (both U>14.50, both *P*<0.004; Fig. 5a). Overall, the lesions reduced the density of cholinergic inputs into the cortex by 50-60%, consistent with the preservation of signal duration-dependent performance in lesioned rats (Fig. 2d,e).

**Figure 4.**
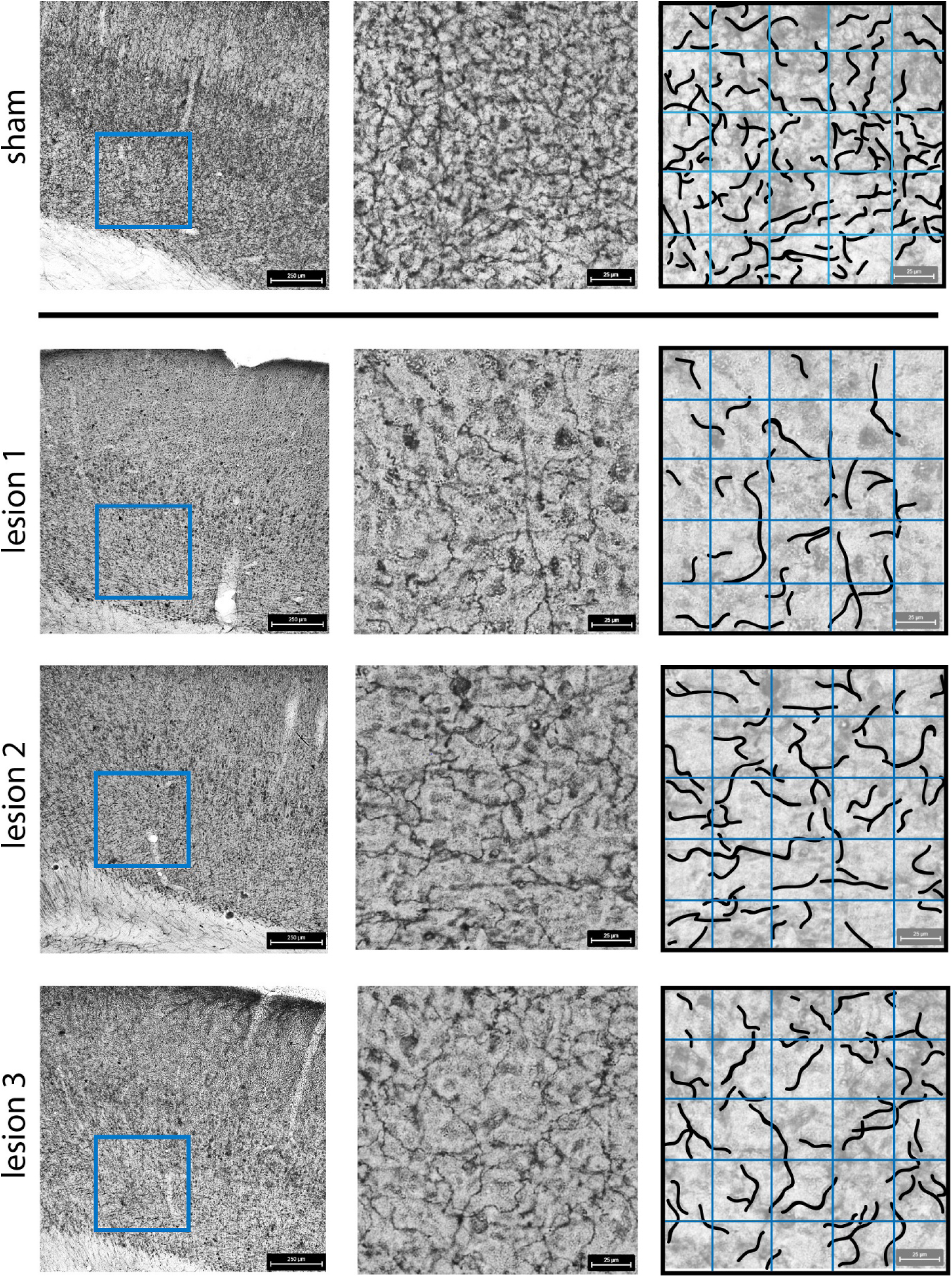
Photomicrographs of AChE-positive fiber stains of the sensorimotor cortex of a sham-lesioned rat (top line) and 3 SAP-lesioned rats (Lesion 1-3). The photographs in the left column (250-µm scale inserted) show the area in layers 5 and 6 of the cortex (blue square) in which AChE-positive fibers were counted at 40× (see magnification in the middle column, 25-µm scale inserted). The right column illustrates the counting of stained fibers using a super-imposed grid method (see Methods for details). For the counting areas shown in the middle column, the counts were (top-to-bottom) 90, 31, 37, and 42.

**Figure 5.**
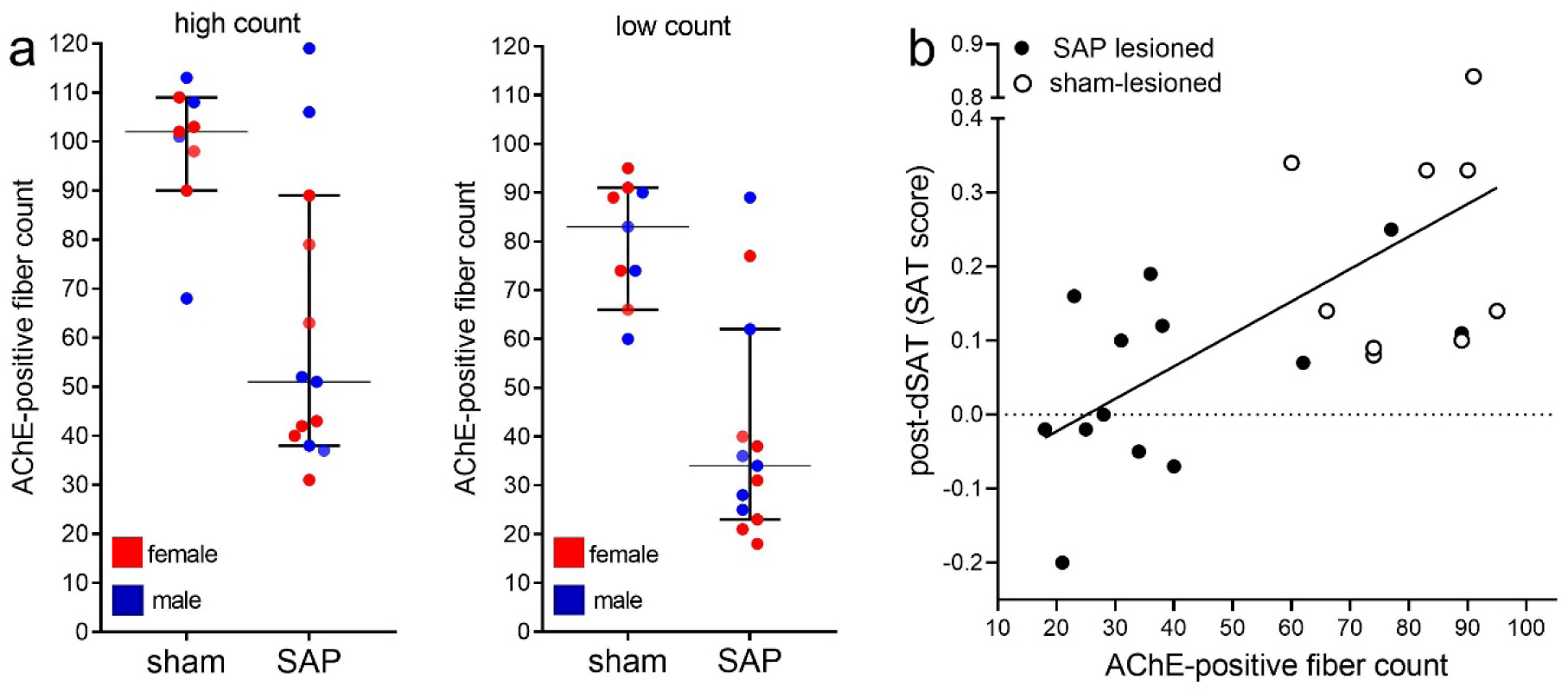
AChE-positive fiber counts were taken from both hemispheres. These counts were highly correlated but also differed significantly. However, for both high and low counts, the effects of the lesions on AChE-positive fiber density were significant (<u>a</u>). Low AChE-fiber counts correlated significantly with the performance of rats (SAT scores) during the post-distractor period (the ordinate depicts data from vehicle-treated rats; note that removal of the outlier – a SAT score of 0.84 - did not abolish this correlation; see Results). Thus, the (residual) density of the cholinergic innervation of the cortex predict the ability of rats to perform following during the task period when demands on attentional control were relatively highest (see Discussion), and when administration of TAK-071 produced significant effects on performance.

Inspection of individual data points in Fig. 5a indicated the counts of three SAP-lesioned rats (2 males, one female) were within the range of the counts from sham-lesioned rats. A *post hoc* exploration of the impact of removing these three rats from the analysis of drug effects on post-dSAT indicated that trends for interactions between the effects of dose, sex, and group, described above, now reached significance.

The (residual) density of the cortical cholinergic input system (low counts) correlated significantly with the recovery of the performance of rats during the post-distractor period (Fig. 5b; SAT scores following vehicle administration versus fiber counts: ρ=0.57, P=0.0053; note that removal of the outlier in Fig. 5b, upper right corner, would not affect this correlation, ρ=0.51, P=0.0015). Likewise, post-dSAT hit rates (from vehicle-treated rats) correlated with AChE-positive fiber counts (ρ=0.43, P=0.043; not shown). Thus, as would be expected, greater cholinergic losses predicted lower performance, specifically during the task period (post-dSAT) when demands on attentional control are relatively high. Although AChE-positive fiber counts did not significantly correlate with measures of drug effects, as indicated above, lower post-dSAT performance predicted greater efficacy of TAK-071.

## DISCUSSION

Removal of 50-60% of the cortical cholinergic input system in rats resulted in significant impairments in SAT performance, specifically hit rates, and in attenuated performance during the post-distractor (post-dSAT) period. Cholinergic losses were highly correlated with post-dSAT performance scores. Administration of TAK-071 (6 days of 0.1 or 0.3 mg/kg; p.o.) neither affected regular SAT performance nor the disrupted performance in the presence of a distractor. However, the drug improved post-dSAT performance (main effect of dose). In rats with relatively lower post-dSAT performance in the absence of drug, administration of the lower dose of TAK-071 yielded relatively higher levels of post-dSAT performance. Moreover, TAK-071 had relatively greater effects in lesioned over sham-operated rats, specifically in terms of enhancing perceptual sensitivity. In addition, a trend for drug-induced improvement of performance in sham-operated females, and a significant drug-induced shift toward a more liberal response bias in females, further suggest, as will be discussed below, that TAK-071 acted by enhancing and rescuing performance in interaction with relatively high demands on attentional control.

### Cholinergic losses and patterns of performance impairments

In this study, cholinergic losses were intended to remain limited to model the losses seen in patients with MCI, early dementia, and PD fallers (see Introduction and Methods for references). Even following more extensive cholinergic losses, impairments in SAT performance do not involve decreases in correct rejections (McGaughy et al., 1996), consistent with the essential role of cholinergic signaling for the detection of cues (Parikh et al., 2007; Howe et al., 2013; Gritton et al., 2016; Howe et al., 2017).

The performance impairments observed in the present experiment were, by design, relatively smaller than those seen in prior studies in rats with more complete cholinergic losses. Furthermore, in the present experiment, the density of cholinergic fibers in cortex correlated with post-dSAT performance (data from vehicle-treated rats), suggesting that relatively limited cholinergic losses most significantly impair performance under conditions that tax attentional control (e.g., Kim and Cave, 1999; Connor et al., 2004). As this type of distractor disrupts performance with considerable efficacy, post-dSAT performance recovery depends particularly closely on cholinergic activity (St Peters et al., 2011), and thus is expected to be attenuated in subjects with cholinergic losses. In humans, such distractors less completely disrupt performance acutely, reflecting their substantially greater attentional control capacity when compared with rodents (Demeter et al., 2008; Demeter et al., 2011). Therefore, in humans, cholinergic losses are expected to be correlated with performance in the presence of the distractor (dSAT) *per se*. This is an important translational qualification of our model and the present data that deserves to be emphasized: While, in rats, the taxation of cholinergic activity and attentional control (for definitions of this theoretical construct see Sarter and Paolone, 2011; Sarter et al., 2016a) cannot be seen during the dSAT and thus necessitates post-dSAT performance data, dSAT performance in humans likely is sufficient to reveal their (attenuated) capacity for cholinergic-attentional control. The findings that resistance to content-rich distractors in PD patients can be attributed to cortical, but not thalamic, cholinergic denervation (Kim et al., 2017b), or that the attentional impact of a genetic sub-capacity variant of cholinergic neurons manifests exclusively in the presence of a distractor (Berry et al., 2014), support for this view.

### Drug effects restricted to post-dSAT

As post-dSAT performance therefore is considered a rat model of dSAT performance in humans, the finding that TAK-071 selectively benefited post-dSAT performance is consistent with the pro-cholinergic nature of the drug effects and has important translational implications. Although the impact of a distractor on the generation of fast cholinergic transients in SAT-performing animals is not known, cholinergic activity monitored by using microdialysis was found to be significantly elevated during dSAT relative to regular SAT performance (St Peters et al., 2011). As elevated extracellular ACh levels collected by microdialysis are hypothesized to reflect integrated transients (Giuliano et al., 2008; Sarter and Lustig, 2019), in subjects attempting to regain above-chance performance in the presence of, or after, the distractor, such a deployment of attentional effort (Sarter et al., 2006) may be mediated via cholinergic transients with greater amplitudes in order to recover cue detection rates (Gritton et al., 2016).

In partially deafferented rats, cholinergic transients likely remain truncated because a reduced number of cholinergic neurons and terminals contribute to the burst of ACh release in cortex. A PAM at M1 receptors would be expected to normalize the postsynaptic consequences of truncated cholinergic transients and therefore to improve the cue detection rates of lesioned rats. The drug-induced increase in *d’* of lesioned rats (Fig. 3e) is consistent with this expectation. It is also noteworthy that, in this analysis, both doses of TAK-071 seemed to increase perceptual sensitivity in lesioned rats even though *post hoc* multiple comparisons attributed the significant interaction between the effects of group and dose on *d’* to the effects of only the lower dose. In contrast, in sham-operated rats, the higher dose of TAK-071 depressed *d’* which, applying our working hypothesis, suggests that the amplification of the postsynaptic effects of cholinergic transients in control rats generally follows a normal distribution: In lesioned rats, truncated cholinergic transients and impaired performance form data points on the left part of a bell-shaped curve and shift upwards following the treatment with TAK-071. In control rats, the amplitudes of cholinergic transients may be near the optimum in the absence of drug, and these data points shift to the right and downwards following further amplification of the postsynaptic cholinergic activity by a M1 PAM.

A similar scenario may also be deduced form our understanding of the postsynaptic, M1-receptor-mediated effects of cholinergic transients. M1-mediated high-frequency oscillations and theta-gamma coupling synchronicity in cortex are necessary to mobilize larger networks involved in cue-guided decision making (Howe et al., 2017). In partially deafferented rats, such gamma oscillations may be robustly suppressed and synchronicity across multiple frequency bands may be disrupted. An M1 PAM would be expected to restore high frequency synchronicity and cross-frequency coupling in deafferented rats while, in intact rats, augmentation of such synchronicity remains without effects or, as illustrated in Fig. 3e, is non-adaptive (see also Rodriguez et al., 2004; Vijayraghavan et al., 2018). These considerations also suggest that effective doses of M1 PAMs for treating patients will need to be carefully determined.

Additional support for interpreting the effects of TAK-071 as mediated by pro-cholinergic mechanisms is based on the results from the analysis of the probabilities for shift-hits. Against our expectation, we found only a trend for cholinergic losses to decrease the probability for shift-hits, perhaps reflecting the presence of relatively limited cholinergic losses. However, consistent with the view that TAK-071 restores the post-synaptic impact of truncated or absent cholinergic activity, the increases in hits produced by TAK-071 (at 0.1 mg/kg) were significant specifically for trials which depend on cholinergic activity, that is, shift-hits, but not for consecutive hits which were previously found not to elicit cholinergic transients (Howe et al., 2013). This finding provides additional insight into the nature of cue detection operations that may benefit from the treatment in TAK-071 in patients. Simple continuous performance tests used to assess attention, where each trial is a cued trial, are less likely to reveal efficacy of TAK-071. Rather, tests which involve shifts between monitoring for cues and cue-oriented behavior, which model real-life attentional situations more closely than monotonous strings of cues (Serences et al., 2005; MacLean et al., 2009; Chun et al., 2011; Henseler et al., 2011), are expected to reveal the therapeutic potential of this M1 PAM.

### “Riskier” response bias favors larger effects of TAK-071

When compared with lesioned males, lesioned female rats generally exhibited a more liberal or a “riskier” response bias, that is, a relatively greater propensity for reporting the presence of cues, along with the costs of such a propensity, a relatively higher false alarm rate. This sex-specific response bias emerged particularly clearly in the dSAT (Fig. 2b,c). As (lesioned) females already exhibited a more liberal bias in the absence of drug, the strong trend for greater efficacy of the drug on the performance of females (Fig. 3c) may have been a function of a further, significant exaggeration of this bias evoked by TAK-071 (Fig. 3f). The M1 PAM is hypothesized to foster the (cholinergic) detection of cues (above), and thus this drug may be relatively more effective in subjects which generally tend to exhibit a bias towards reporting the presence of cues.

It is difficult to translate these findings to direct predictions of drug effects on sex-specific response biases in humans, once again because of their inordinately greater capacity for distractor filtering and for modulating cue processing in the presence of distractors (Weissman et al., 2002; Demeter et al., 2008; Demeter et al., 2016), Moreover, in response to shifts in trial outcome, humans shift their biases more accurately and sensitively than rats (Echevarria et al., 2005; Lynn and Barrett, 2014). Thus, the role of sex-specific response biases in modulating drug effects in rats may not translate into the role of sex-specific biases in humans but, more generally, may indicate that the efficacy of TAK-071 in clinical populations may be a function of the subjects’ response biases and the sensitivity of response biases to task manipulations. If response biases remain relatively insensitive to task manipulations, as was the case in the male rats in the present experiment, less robust effects of TAK-071 may be expected.

## Conclusions: Translational significance

Because of the paucity of potential therapeutic compounds which have been assessed both in the present model and patients, the evaluation of the potential translational significance of the present findings relies on the construct validity of the present experimental approach. The limited degree of cholinergic depletion in rats, designed to model specific patient groups (above), the use of a highly translational SAT task (Demeter et al., 2008; Demeter et al., 2011; Demeter et al., 2013; Demeter et al., 2016; Demeter and Woldorff, 2016), and the close relationship between cholinergic transient signaling and performance in this task (references above), are characteristics which indicate the construct validity of the overall neuro-behavioral paradigm. Furthermore, the orderly nature of the present neurobehavioral data (e.g., highly significant correlations between AChE-fiber density and post-dSAT performance, and between drug effects on post-SAT performance and performance in the absence of drugs) may be considered a foundation for suggesting efficacy in patients. The present data suggest that TAK-071 improves the attentional performance in patients with limited losses of the forebrain cholinergic system, including patients with MCI and early dementia (Mesulam et al., 2004; Ray et al., 2015) and, perhaps most likely, in PD patients with cholinergic losses concomitant to their dopamine losses (Bohnen et al., 2009; Kim et al., 2017b). Importantly, drug effects may be revealed primarily in situations characterized by demands on attentional control (Sarter et al., 2016a) and, as discussed above, drug effects may interact with the presence of relatively more liberal and more flexible response biases. In PD patients, the pro-cholinergic and pro-attentional effects of this M1 PAM may also manifest in terms of improved gait and balance control and a reduction in fall rates.

## Author contributions

AK and AKC conducted the experiments. AK and KBP analyzed the data and conducted the histological analyses. MS designed the experiment. AK, KBP and MS wrote the paper.

## Acknowledgements

We thank Dr. Arthur Simen (Takeda, Cambridge, MA) for numerous discussions of this project and Dr. Cindy Lustig (University of Michigan) for comments on a draft of the paper. AKC is now at PsychoGenics (NJ).

